# Cell envelope structural and functional contributions to antibiotic resistance in *Burkholderia cenocepacia*

**DOI:** 10.1101/2024.01.03.574096

**Authors:** Andrew M Hogan, Anna Motnenko, A S M Zisanur Rahman, Silvia T Cardona

## Abstract

Antibiotic activity is limited by the physical construction of the Gram-negative cell envelope. Species of the *Burkholderia cepacia* complex (Bcc) are known as intrinsically multidrug-resistant opportunistic pathogens with low permeability cell envelopes. Here, we re-examined a previously performed chemical-genetic screen of barcoded transposon mutants in *B. cenocepacia* K56-2, focusing on cell envelope structural and functional processes. We identified structures mechanistically important for resistance to singular and multiple antibiotic classes. For example, the polymeric O-antigen was important for resistance to cationic antibiotics, while defects in peptidoglycan precursor synthesis specifically increased susceptibility to cycloserine and revealed a new putative amino acid racemase. Susceptibility to novobiocin, avibactam, and the LpxC inhibitor, PF-04753299, was linked to the BpeAB-OprB efflux pump, suggesting these drugs are substrates for this pump in *B. cenocepacia*. Additionally, disruption of the periplasmic disulfide bond formation system caused pleiotropic defects on outer membrane integrity and β-lactamase activity. Our findings highlight the layering of resistance mechanisms in the structure and function of the cell envelope. Consequently, we point out processes that can be targeted for developing antibiotic potentiators.

**Importance:** The Gram-negative cell envelope is a double-layered physical barrier that protects cells from extracellular stressors, such as antibiotics. The *Burkholderia* cell envelope is known to contain additional modifications that reduce permeability. We investigated *Burkholderia* cell envelope factors contributing to antibiotic resistance from a genome-wide view by re-examining data from a transposon mutant library exposed to an antibiotic panel. We identified susceptible phenotypes for defects in structures and functions in the outer membrane, periplasm, and cytoplasm. Overall, we show that resistance linked to the cell envelope is multifaceted and provides new targets for the development of antibiotic potentiators.

## Introduction

The Gram-negative cell envelope plays a significant role in antibiotic resistance (1), which causes millions of deaths each year (2). In addition to imparting cells with mechanical integrity, shape, and sensory functions, the Gram-negative cell envelope is a selectively permeable surface for solute exchange between the cell and the environment. Its functions are controlled by the composition of the inner and outer membranes (3, 4) and the thin layer of peptidoglycan located in the periplasm (5). The inner membrane contains many specific transporter proteins embedded in a phospholipid bilayer. The outer membrane is a highly asymmetric bilayer with an outer leaflet primarily composed of lipopolysaccharide (LPS) with phospholipids on the inner leaflet (6). While the LPS dramatically reduces the permeability of the lipid portion of the outer membrane (7), water-filled porins that stud the outer membrane generally allow small hydrophilic compounds (less than 600 Da) to diffuse into the periplasm (8). On the other hand, the peptidoglycan matrix is a dynamic structure synthesized by large protein complexes – the elongasome and divisome – and “trimmed” by a variety of lytic enzymes (e.g. transglycosylases and endopeptidases) (9). Peptidoglycan turnover by these lytic enzymes occurs even in the stationary phase (10). While not directly involved with antibiotic permeability, peptidoglycan matrix integrity is linked to regulatory circuits that sense cell wall stress and respond by upregulating β-lactamases, chaperones, and alternative genes involved in matrix synthesis (5). Thus, the peptidoglycan matrix plays an important role in cell envelope-related antibiotic susceptibility and stress responses, apart from its direct targeting by β-lactams.

Gaps in our knowledge of the factors controlling cell envelope permeability and integrity are often cited in the failure of antibiotic discovery platforms (11, 12). A set of predictive “rules” for porin-mediated entry into *E. coli* (13) and *Pseudomonas aeruginosa* (14, 15) have been described. However, the overall challenge of antibiotic entry into the cytoplasm remains due to opposing permeability preferences from the outer and inner membranes (16) and differences in cell envelope construction between bacterial pathogens (17, 18).

Members of the Gram-negative *Burkholderia cepacia* complex (Bcc) cause infections in people with cystic fibrosis and are notorious for their high levels of resistance to multiple antibiotic classes (19). Bcc bacteria contain a variety of modifications in their cell envelope that reduce antibiotic activity. In the LPS core oligosaccharide, D-*glycero*-α-D-*talo*-octulosonic acid (Ko) is commonly found instead of 3-deoxy-D-*manno*-octulosonic acid (Kdo) and is also functionalized with 4-amino-4-deoxy-L-arabinose (Ara4N) (20). The addition of these positively charged Ara4N residues is essential for efficient LPS export to the outer membrane and also contributes to high-level cationic peptide resistance (21, 22). The major porins are also thought to have a narrow channel diameter, limiting small molecule diffusion rates to approximately 10-fold less than *E. coli* and similar to rates determined for *P. aeruginosa* (23). To counter the entry of antibiotics, Bcc species have a suite of broad-spectrum efflux pumps (24). Aside from resistance due to the major β-lactamases (25, 26), the contributions of peptidoglycan biosynthesis and recycling factors to antibiotic susceptibility in the Bcc are unclear.

We previously constructed a barcoded transposon mutant library in the Bcc member *B. cenocepacia* K56-2 and exposed it to diverse cell envelope-targeting antibiotics (26). Strain K56-2 was chosen as it is a highly virulent, multidrug-resistant clinical isolate from the ET12 epidemic lineage (27). Barcode sequencing (BarSeq) was used to uncover connections of cell envelope structural elements to β-lactam susceptibility (26). Here, we report additional findings from the screening data, focusing on cell envelope structural and functional components. We identified new roles in *B. cenocepacia* antibiotic resistance for the periplasmic disulfide bond pathway, peptidoglycan recycling, cell division accessories, specific porins, and additional substrates for the important BpeAB-OprB efflux pump. We expand how the cell envelope acts as an antibiotic barrier and expose weaknesses that may be exploited as targets to increase antibiotic activity.

## Results

To profile cell envelope-related antibiotic resistance in *B. cenocepacia* K56-2, we previously selected 22 antibiotics targeting the cell envelope (Table S1) and used them at subinhibitory concentrations to challenge a barcoded transposon mutant library (26). After quantification of fitness with barcode sequencing (BarSeq), we selected several genes with negative fitness effects (i.e. disruption increased antibiotic susceptibility) that are connected to cell envelope structure and integrity for further investigation.

### LPS, lipid A modifications, and the RpoE regulon confer resistance to cationic antibiotics

As part of their mechanisms of action, the cationic antibiotics chlorhexidine (CHX) and polymyxin B (PMB) bind to and disrupt bacterial membranes. In *E. coli,* PMB binds the lipid A moiety of LPS in the inner and outer membranes, resulting in cell lysis (28). CHX is reported to collapse the membrane potential and preferentially disrupt the outer membrane (29). In other Gram-negatives, such as *Salmonella enterica* and *P. aeruginosa*, cationic antibiotic resistance is inducible (30); however, *Burkholderia* species are intrinsically resistant to these agents, mediated both by physical protection of the membrane and by adaptive responses (31). The *Burkholderia* O-antigen, LPS core, and RpoE envelope stress regulon are important for PMB resistance (31, 32).

We leveraged the BarSeq data for a deeper look into cationic antibiotic resistance in *Burkholderia*. We identified genes important for PMB and CHX fitness associated with O-antigen synthesis, LPS core synthesis, and lipid A synthesis (26). To dissect the importance of the O-antigen versus LPS core, we constructed an unmarked deletion mutant in *wbiI*, encoding an epimerase/dehydratase required for O-antigen synthesis. As expected, this mutant lacked polymeric O-antigen and could be complemented *in trans* (Figure S1). We also used a previously constructed unmarked deletion mutant in *hldD* (26), encoding an epimerase required for LPS core synthesis, which lacks both polymeric O-antigen and a complete LPS core. Both the Δ*wbiI* and Δ*hldD* mutants were markedly more susceptible than wild-type to PMB, colistin (COL) and CHX; however, the Δ*wbiI* mutant was generally less susceptible than the Δ*hldD* mutant, and the increase in CHX susceptibility was only modest for both mutants (Figure 1A). The stepwise change in cationic antibiotic susceptibility for mutants in the O-antigen versus LPS core suggests each has a separate but complementary role in physically protecting the cell.

**Figure 1.**
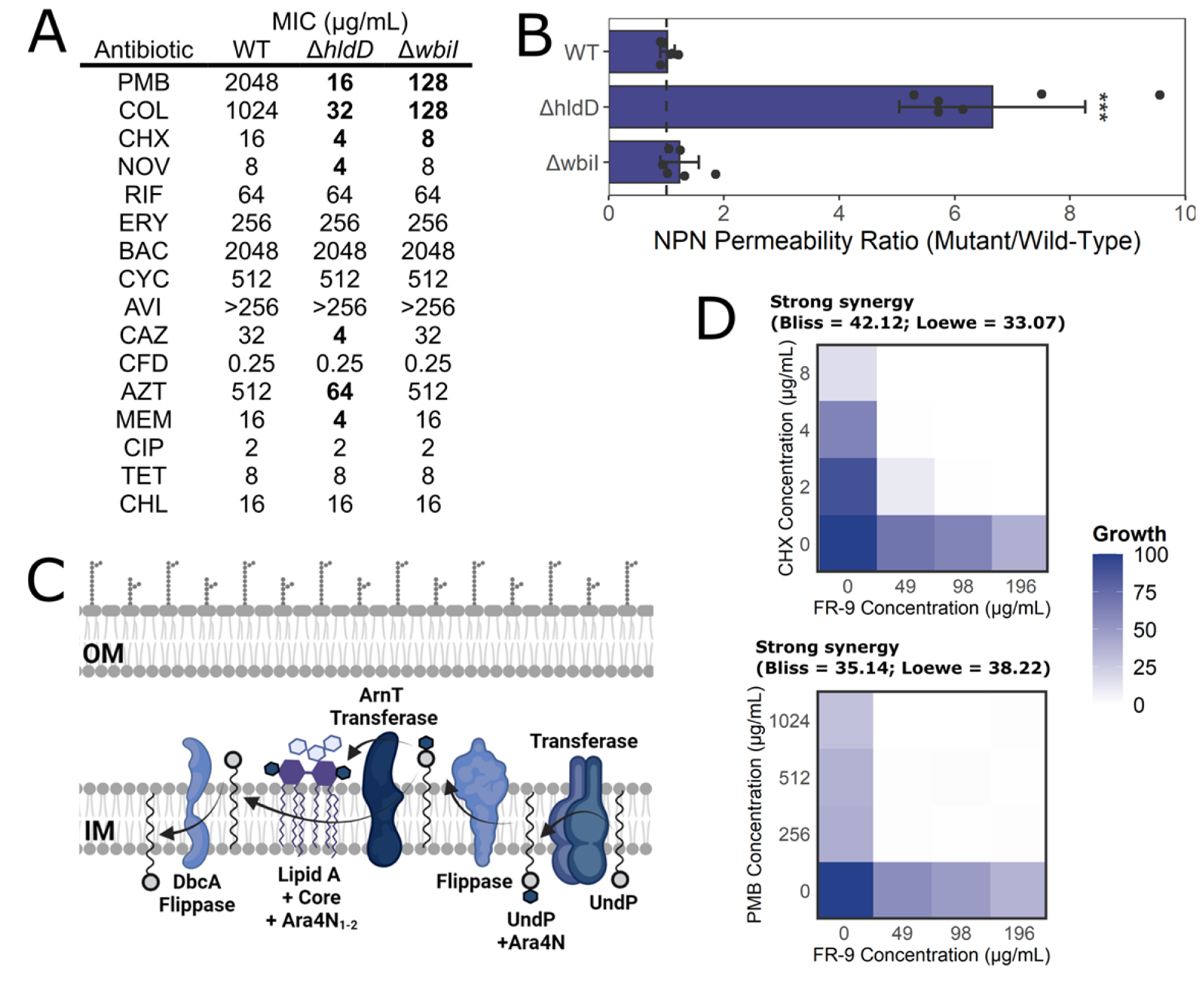
Defects in the O-antigen, LPS core, lipid A modifications, and RpoE regulon increase susceptibility to cationic antibiotics. A) MIC values of K56-2 and the Δ*hldD* and Δ*wbiI* mutants. Values are medians of three biological replicates, with bold indicating at least a 2-fold change from WT. B) Ratios of NPN fluorescence of the deletion mutants to the fluorescence of the wild-type control. Error bars represent means ± SD of six replicates. Significance was determined by 1-way ANOVA with Dunnett’s *post hoc* test to the wild-type control. *** P < 0.001. The dashed line indicates an NPN fluorescence ratio of 1. C) Diagram of the membrane-associated steps of the Ara4N LPS modification, inferred from homology and experimental evidence in *Burkholderia*. D) Mini checkboard assays of FR-9 interaction with CHX and PMB. Values are normalized to the OD_600_ of growth without antibiotics. On top of each plot are the interaction values and interpretation from SynergyFinder.

As LPS is an important structural component of the outer membrane, we next addressed whether the Δ*wbiI* and Δ*hldD* mutants also had membrane permeability defects. We incubated cells with 1-*N*-phenylnaphthylamine (NPN). This hydrophobic small molecule becomes fluorescent when dissolved in the lipids of the outer membrane (33, 34), a space from which NPN is normally excluded in cells with intact membranes. Compared to the wild-type strain, only the Δ*hldD* mutant displayed significantly increased outer membrane permeability (Figure 1B). This mutant contains a complete LPS core but not an O-antigen (ref). However, while the Δ*hldD* mutant was more susceptible to the β-lactams aztreonam (AZT), ceftazidime (CAZ), and meropenem (MEM), this mutant was not more susceptible to the large scaffold antibiotics erythromycin (ERY), novobiocin (NOV), and rifampicin (RIF) (Figure 1A), as would be expected for a mutant with a compromised outer membrane. Additionally, the magnitude of BarSeq gene fitness of *hldD* was not correlated with antibiotic molecular weight (P = 0.27; Figure S2), suggesting the permeability defect of the Δ*hldD* mutant is only moderate. On the other hand, the Δ*wbiI* mutant did not have increased susceptibility to either the β-lactams of large scaffold antibiotics.

Lipid A modification with 4-deoxy-4-amino-arabinose (Ara4N) provides cationic antibiotic resistance by reducing the net negative charge of LPS (35). Undecaprenyl phosphate (UndP) is known to be the carrier for Ara4N (36) (Figure 1C). The *E. coli dedA* encodes an UndP flippase, which is critical for UndP recycling (37–39). We previously found that disruption of *dbcA*, encoding a homologue of the *E. coli dedA* (bi-directional best-hit BLAST), increased susceptibility to PMB and CHX (26). Supporting the connection between UndP and the Ara4N modification, we found that inhibition of UndP synthesis with the fosmidomycin analogue FR-900098 (FR-9) strongly synergised with both PMB and CHX (Figure 1D).

### Novobiocin, avibactam, and the LpxC inhibitor PF-04753299 are substrates for the BpeAB-OprB efflux pump and specific porins

Bacteria possess a variety of efflux pumps that are associated with a diverse range of functions, including the removal of toxic compounds such as antibiotics. However, many of the antibiotics we used are not known to be substrates for *Burkholderia* efflux pumps. *B. cenocepacia* K56-2 encodes 16 tripartite RND efflux pumps, of which several are important for antibiotic resistance (40). We hypothesized that transposon insertions in genes encoding efflux pumps would decrease fitness in the presence of each pump’s range of substrates.

From the BarSeq experiment, we observed that disruption of the K562_RS30055 PACE family efflux pump decreased fitness in the presence of multiple antibiotics. In contrast, disruption of the K562_RS10630 EmrE pump homologue only decreased fitness with CHX (Figure 2A). Additionally, we found that disruptions in components of the BpeAB-OprB efflux pump (also called RND-4, a homologue of the MexAB-OprM pump in *P. aeruginosa*) increased susceptibility to NOV, avibactam (AVI), AVI/CAZ, PF-04753299 (PF-04) and CHIR-090 (CHIR) (Figure 2A). Enhanced susceptibility was only seen for disruptions in *bpeA* and *bpeB* but not for *oprB*, suggesting that the *bpeAB* complex may also form functional associations with other outer membrane factors. NOV, AVI, PF-04, and CHIR are not known to be effluxed by BpeAB-OprB in *B. cenocepacia*. To verify these findings, we used CRISPRi (41) for rhamnose-inducible knockdown of the *bpeABoprB* operon. Knockdown was validated by qRT-PCR (Table S2). Knockdown of this pump did not cause a growth defect in the absence of antibiotics (Fig S3A). Silencing by CRISPRi was validated by a 2-fold reduction in MIC of the known substrate chloramphenicol (CHL) (Figure 2B). Upon dCas9 induction, susceptibility to NOV, PF-04, AVI, and AVI/CAZ moderately increased, but not to CAZ only (Figure 2B). Additionally, we overexpressed the *bpeABoprB* operon from a multicopy plasmid and, with careful titration of rhamnose to prevent a growth defect (Figure S3D), we observed moderately reduced susceptibility to CHL, NOV, PF-04, AVI, and AVI/CAZ (Figure 2B). We thus conclude that NOV, AVI, and PF-04 are also substrates for the BpeAB-OprB efflux pump in *B. cenocepacia*.

**Figure 2.**
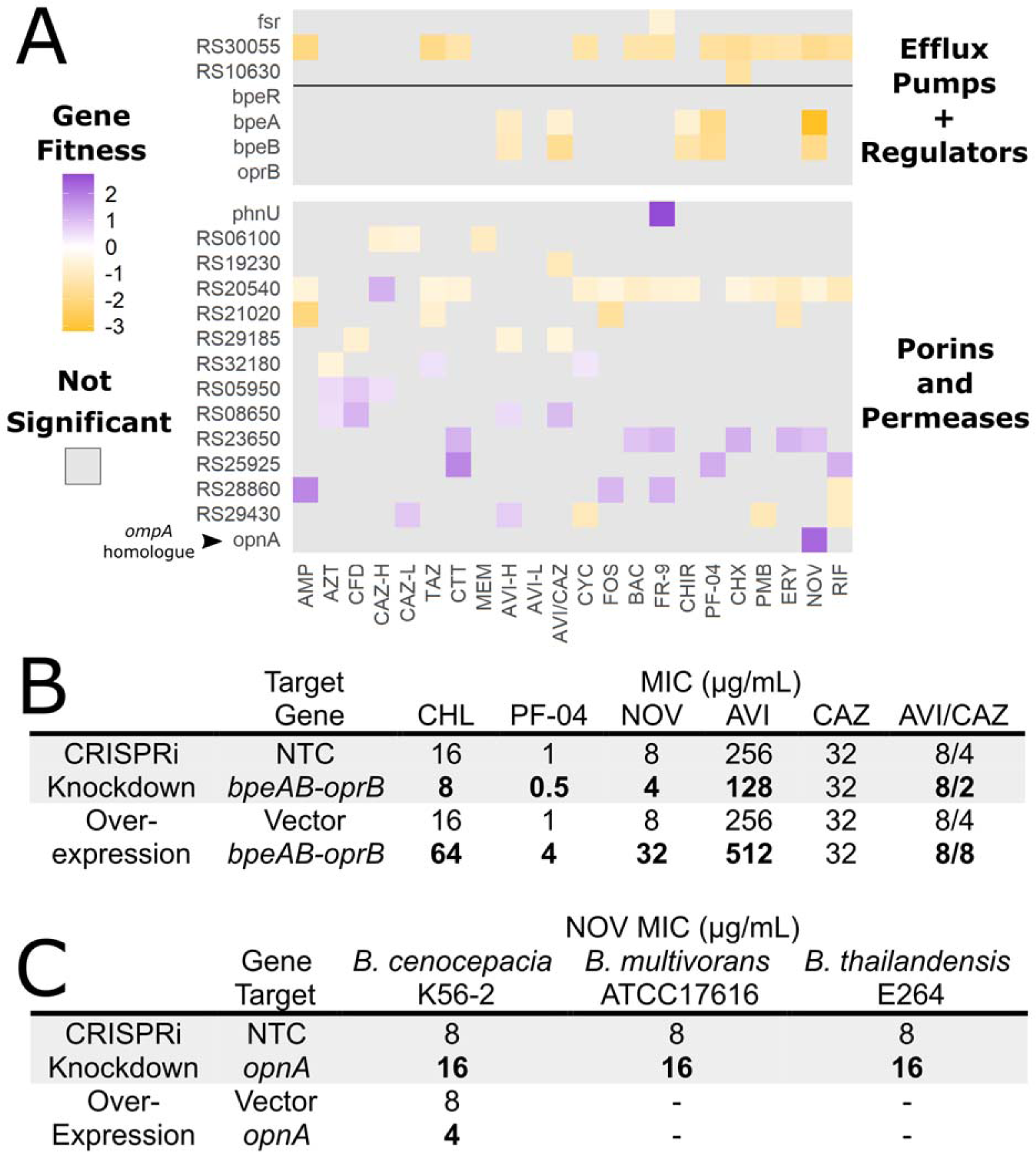
Disruptions in efflux pumps, porins, and permeases alter antibiotic susceptibility. A) BarSeq gene fitness scores relative to DMSO control of the genes associated with efflux pumps, porins, and permeases. The K562_# locus tags are given for genes without assigned names. Grey indicates the interactions were not significant (P > 0.05). B) MIC values of CRISPRi and overexpression mutants of the *bpeABoprB* operon in K56-2. C) MIC values of CRISPRi and overexpression mutants of *opnA* (K562_RS14130) in *Burkholderia* species. NTC = non-targeting sgRNA control. MIC values are medians of three biological replicates, with bold indicating at least a 2-fold change from the control when rhamnose was added (0.5% for CRISPRi mutants; 0.005% for overexpression).

K56-2 encodes over 130 putatively annotated porins and permeases, and few have been characterised by their physiological importance. We observed that disruption of K562_RS29850, encoding a homologue of the PhnU 2-aminoethylphosphonate ABC-type permease, resulted in markedly reduced susceptibility exclusively in the presence of FR-9 (Figure 2A). The structural similarity between 2-aminoethylphosphonate and FR-9 suggests that FR-9 may also be taken into K56-2 via this permease. We found that other porins were important for fitness in multiple conditions (e.g. K562_RS21020 and K562_RS20540) (Figure 2A). Some porins, such as OmpA and OmpC in *E. coli*, are important for maintaining membrane integrity (42), which may explain why the lack of porins may reduce fitness in the presence of antibiotics. On the other hand, we observed several porins that, when disrupted, increased fitness in at least one condition (e.g. K562_RS23650 and K562_RS25925) (Figure 2A). No porins were associated with fitness in all conditions, not even for all β-lactams, which suggests an intricate network of minor variations in porin specificity and partial functional redundancy.

Of note, we observed that disruption of K562_RS14130, a homologue of *E. coli* K-12 *ompA* (53.4% identity, top hit by reciprocal best-hit BLAST), exclusively increased fitness in the presence of NOV (Figure 2A). In *E. coli*, a deletion of *ompA* does not affect susceptibility to NOV (42). Phylogenetics revealed that the porin encoded by K562_RS14130 is conserved with greater than 60% amino acid identity among many *Burkholderiaceae* genera (Figure S4A). The presence of this porin also appears to coincide with low MIC values of NOV, but only for species of *Burkholderia* (Figure S4B). In K56-2, *B. multivorans* ATCC 17616 and *B. thailandensis* E264, CRISPRi repression of the K562_RS14130 homologue modestly reduced susceptibility to NOV (Figure 2C), but not other antibiotics (Figure S4C). Additionally, overexpression of K562_RS14130 in K56-2 increased susceptibility to NOV (Figure 2C). Together, our findings support a different role of K562_RS14130 from *E. coli ompA,* and we propose it to be annotated as *opnA* for <u>O</u>uter membrane <u>P</u>rotein, <u>N</u>ovobiocin entry <u>A</u>.

### The DSB system has a pleiotropic effect on antibiotic resistance

The folding and stability of secreted proteins are aided by disulfide bond (DSB) formation via the periplasmic DSB system. In *E. coli*, this system is encoded by four genes organized in two pathways, the DsbAB pathway and the DsbCD pathway, required for efficient disulfide bond formation and isomerization, respectively, in periplasmic proteins (Figure 3A) (43). Reports from the pre-genomic era have associated DsbAB in *Burkholderia* with effects on protease activity, motility, and resistance to multiple antibiotics (44, 45). We found that transposon disruptions in *dsbA* and *dsbB* resulted in large changes in susceptibility to many β-lactams and large scaffold antibiotics. In contrast, disruptions in *dsbC* and *dsbD* did little to alter the susceptibility profile (Figure 3B).

**Figure 3.**
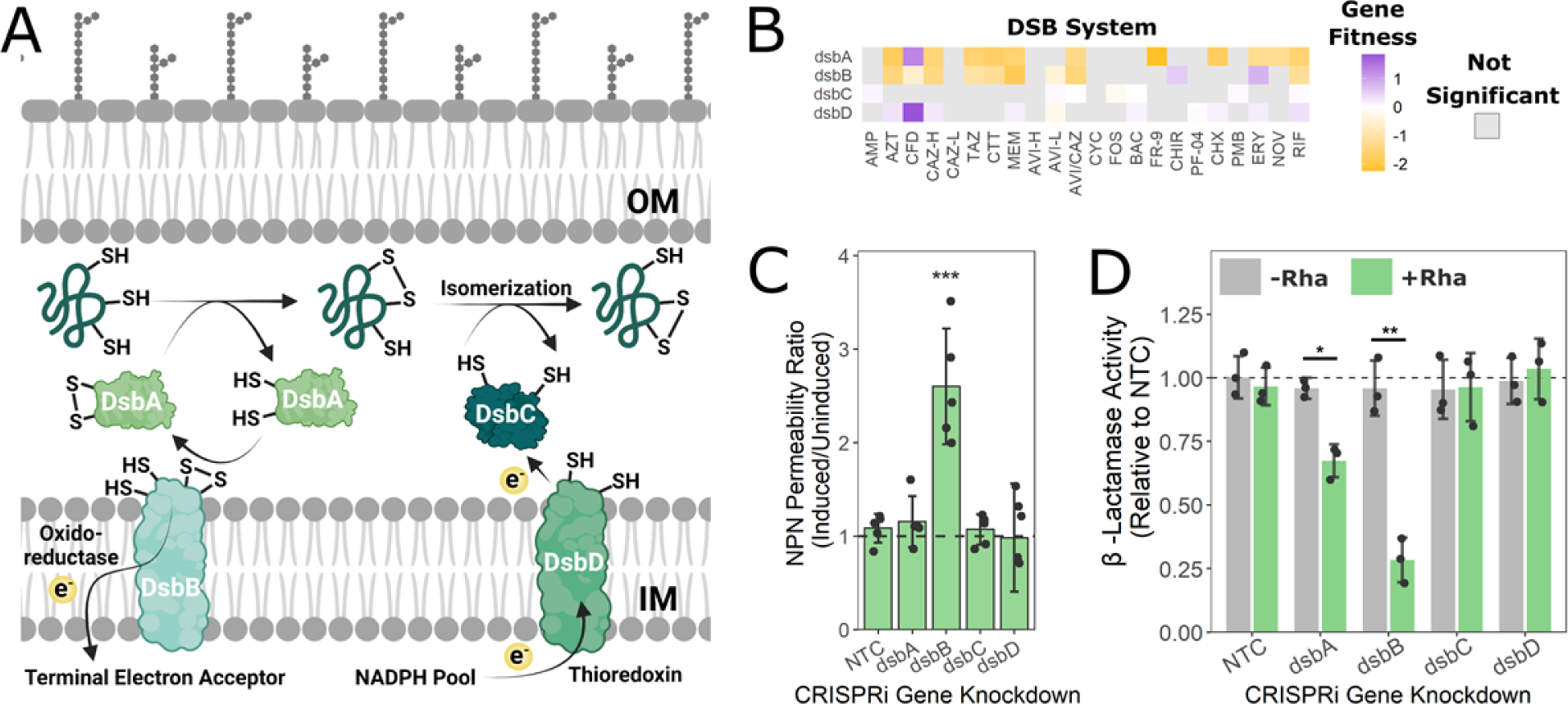
Defects in the DSB system increase outer membrane permeability and decrease β-lactamase activity, resulting in susceptibility to multiple antibiotics. A) Diagram of periplasmic disulfide bond formation in nascent proteins by DsbAB and isomerization by DsbCD. B) BarSeq gene fitness scores relative to the DMSO control of the genes in the DSB system. Grey indicates the interactions were not significant (P > 0.05). C) Ratios of NPN fluorescence of the CRISPRi mutants in inducing (0.5% rhamnose for *dsbA*, *dsbC*, and *dsbD*; 0.05% rhamnose for *dsbB*) vs. uninducing conditions (0% rhamnose). Values represent means ± SD of five biological replicates. Significance was determined by 1-way ANOVA with Dunnett’s *post hoc* test to the wild-type control. *** P < 0.001. The dashed line indicates an NPN fluorescence ratio of 1. D) β-lactamase activity of CRISPRi mutants in the DSB system assayed by nitrocefin hydrolysis of lysates of cells grown with or without rhamnose induction. Values represent means ± SD of five biological replicates. The dashed line indicates the activity of the control sgRNA (NTC). Significance was determined by an unpaired two-tailed t-test to the NTC grown without rhamnose using Bonferroni’s correction. * P < 0.05; ** P < 0.01.

Hundreds of extracellular proteins in a typical cell may be oxidized by the DSB system (46, 47). By screening predicted secreted proteins (48) in K56-2 for at least two cysteine residues, we identified 615 putative DSB system substrates (Table S3). With a more stringent threshold of an even number of cysteine residues (46), 416 proteins were identified (Table S3). As expected, these putative substrates are enriched in cell envelope-associated processes (Table S4). As envelope structural proteins (e.g. LptD) were among the putative DSB system substrates, this raised the possibility that disruptions in *dsb* genes affected envelope integrity. However, only knockdown of *dsbB* significantly increased outer membrane permeability (Figure 3C). Additionally, β-lactamases have previously been implicated as substrates of DsbA in *P. aeruginosa* (49). As PenB, the dominant β-lactamase in K56-2 (26), was also identified as a putative DSB system substrate (Table S3), we determined β-lactamase activity in targeted CRISPRi mutants using a nitrocefin hydrolysis assay. Knockdown of *dsbA* and *dsbB* reduced β-lactamase activity by ∼30% and ∼75%, respectively, while knockdown of *dsbC* and *dsbD* showed no difference from the control (Figure 3D). Together, we demonstrate that the pleiotropic effect of the DSB system on outer membrane permeability and periplasmic β-lactamase activity in K56-2 is important for resistance to multiple classes of antibiotics.

### Disruptions in peptidoglycan synthesis and recycling genes cause susceptibility to β-lactams

β-lactams remain one of the few options to treat Bcc infections. We thus decided to examine the genome-wide β-lactam susceptibility profiles with a focus on their target: peptidoglycan metabolism. As the primary target of the β-lactams are penicillin-binding proteins (PBPs), we expected that disruption of specific PBPs may expose weaknesses in peptidoglycan matrix biosynthetic processes. We observed that disruption of only two PBP-encoding genes altered β-lactam susceptibility: K562_RS05010 (encoding a homologue of the DacB DD-carboxypeptidase, also called PBP4) and K562_RS01445 (encoding a homologue of *E. coli mrcA/pbp1a* we are referring to as *mrcA2*) (Figure 4A). Very low fitness in the presence of AVI and MEM was exclusive to the disruption in *dacB*, while impaired fitness in the presence of AZT, tazobactam (TAZ), cycloserine (CYC), fosfomycin (FOS), and cefiderocol (CFD) was exclusive to the disruption in *mrcA2*. Reduced fitness in cefotetan (CTT), CAZ-H, and AVI/CAZ were common to both disruptions in *dacB* and *mrcA2* (Figure 4A). As disruption of *dacB* resulted in very strong fitness effects in the BarSeq conditions, we targeted it for CRISPRi knockdown. In the absence of antibiotics, the mutant displayed no growth defect (Figure S3A). However, silencing *dacB* greatly increased susceptibility to MEM and AVI (Figure 4B), the two strongest interactions for *dacB* in the BarSeq experiment.

**Figure 4.**
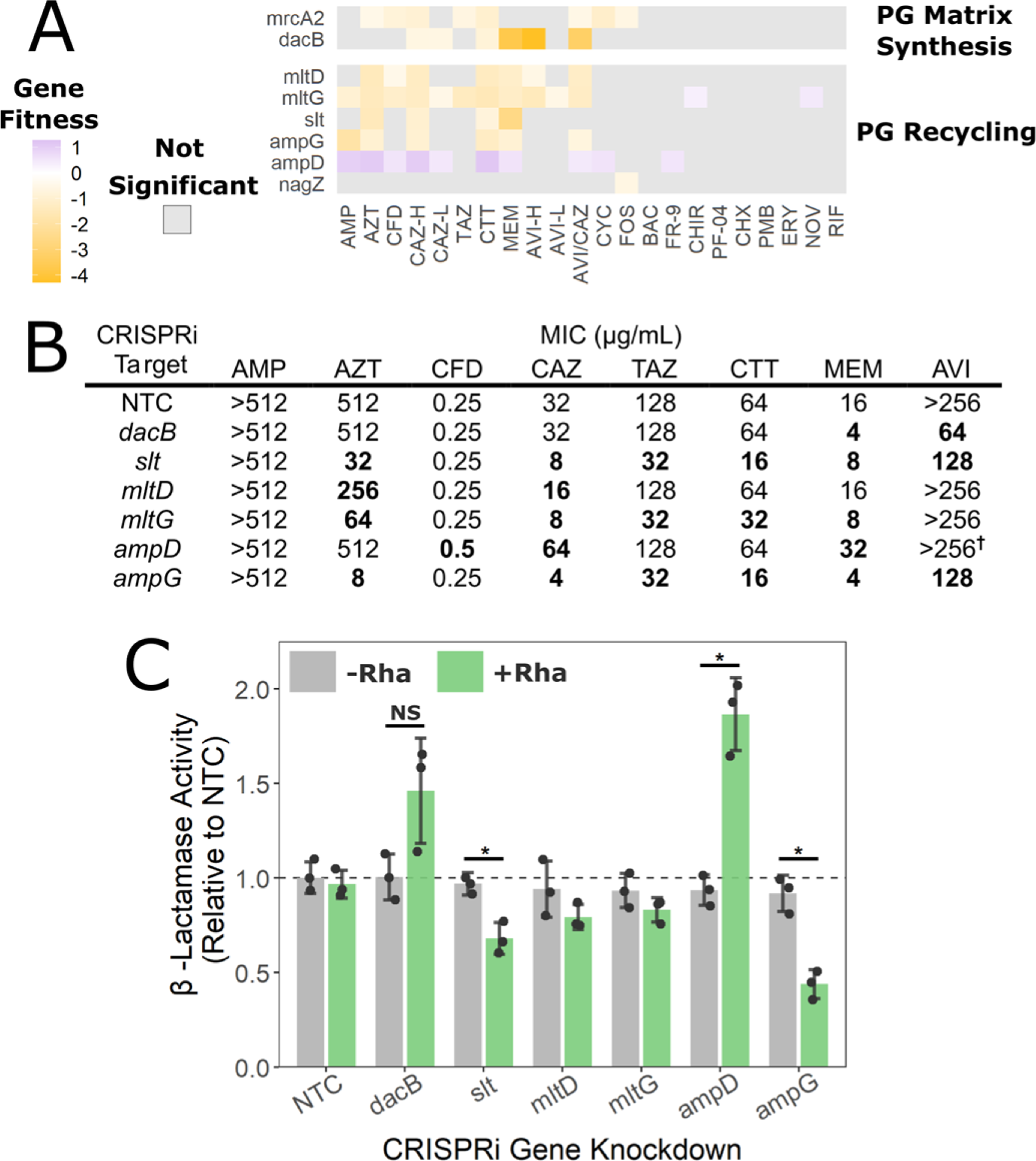
Disrupting peptidoglycan matrix synthesis and recycling genes increases β-lactam susceptibility. A) BarSeq gene fitness scores relative to DMSO control of the genes encoding peptidoglycan recycling and matrix synthesis processes. Grey indicates the interactions were not significant (P > 0.05). B) MIC values of K56-2 CRISPRi mutants harbouring plasmids expressing a non-targeting sgRNA control (NTC) or a sgRNA targeting the indicated genes. Values shown are MIC values (medians of three biological replicates) in the presence of rhamnose induction of CRISPRi, with bold indicating at least a 2-fold change from the NTC. † MIC value did not change, but growth at subinhibitory concentrations was substantially higher. C) β-lactamase activity of indicated CRISPRi mutants assayed by nitrocefin hydrolysis of cell lysates. Values represent means ± SD of three biological replicates. The dashed line indicates the activity of the control sgRNA (NTC). Significance was determined by an unpaired two-tailed t-test to the NTC grown without rhamnose using Bonferroni’s correction. * P < 0.05; NS = not significant (P > 0.05).

Notably, the findings from the BarSeq experiment are in contrast with results obtained in *P. aeruginosa* (50), *Yersinia enterocolitica* (51), and *Aeromonas hydrophila* (52), which show that inactivation of DacB causes β-lactam resistance. In *P. aeruginosa*, the disruption of DacB leads to activation of AmpR and the CreBC/BlrAB two-component system, resulting in AmpC overexpression (50). Disruption of the K56-2 CreBC/BlrAB homologues (K562_RS27235-40) did not affect fitness in any condition in the BarSeq experiment (26). Additionally, we did not see a significant change in β-lactamase activity when we knocked down *dacB* (Figure 4C). Overall, we suggest that inactivation of DacB and the consequences of its inactivation may play a different physiological role in *B. cenocepacia*.

Peptidoglycan recycling co-occurs with peptidoglycan synthesis, and changes in the balance alter β-lactam resistance. In *E. coli,* various endopeptidases, lytic transglycosylases, and transporters are required for peptidoglycan recycling (53). However, very little is known about peptidoglycan recycling in *Burkholderia*. Analysis of the BarSeq data for homologues of genes associated with peptidoglycan recycling revealed several as important for fitness in the presence of β-lactams (Figure 4A). Disruptions in *slt* (encoding a soluble lytic transglycosylase) and in *mltD* and *mltG* (each encoding a membrane-bound lytic transglycosylase) all resulted in susceptibility to many β-lactams, but not the early peptidoglycan synthesis inhibitors CYC and FOS (Figure 4A). To validate our findings from the BarSeq experiment, we silenced *slt, mltD*, and *mltG* with CRISPRi, which variably increased susceptibility to AZT, CAZ, CTT, MEM, TAZ, and AVI (Figure 4B). Knockdown of *slt* was associated with the greatest changes in susceptibility, which may be explained by an approximate 35% reduction in β-lactamase activity (Figure 4C). Neither knockdown of *mltD* nor *mltG* significantly affected β-lactamase activity. While other endopeptidases and transglycosylases are encoded in K56-2 in addition to *slt*, *mltD*, and *mltG*, they did not contribute to fitness in the conditions used for the BarSeq experiment, perhaps due to partial functional redundancy, as recently demonstrated in *V. cholerae* (54).

The balance of various peptidoglycan degradation products has important implications for β-lactamase expression. In *Citrobacteri freundii*, β-lactam exposure causes lytic enzymes to liberate GlcNAc-anhydroMurNAc-pentapeptide fragments, which enter the cytoplasm through the AmpG permease (55) and then are processed by NagZ into anhydroMurNAc-pentapeptide, then into anhydroMurNAc by AmpD (56). Both the substrate and product of NagZ, but not the product of AmpD, activate the expression of β-lactamase genes (57, 58); thus, AmpD activity reduces β-lactamase expression. In *B. cenocepacia,* while mutations in *ampD* are known to confer β-lactamase resistance (59), the contributions of AmpG have not been experimentally characterised. In the BarSeq experiment, disruption of *ampG* increased susceptibility to many β-lactams. In contrast, disruption of *ampD* reduced susceptibility to many β-lactams and no effect was observed with disruption of *nagZ* (Figure 4A). Upon silencing of *ampD* with CRISPRi, susceptibility was reduced to CAZ, CFD, and MEM, while silencing of *ampG* increased susceptibility to AZT, CAZ, TAZ, CTT, and MEM (Figure 4B). Supporting these trends, a knockdown of *ampD* nearly doubled β-lactamase activity, while the β-lactamase activity of the *ampG* knockdown was almost halved (Figure 4C). Neither knockdown affected growth in the absence of antibiotics (Figure S3A). Thus, our findings support the role of AmpG as a permease for peptidoglycan degradation products and of AmpD as reducing the accumulation of anhydroMurNAc-pentapeptides in K56-2, and both being important for β-lactamase regulation.

### Peptidoglycan precursor synthesis modifies susceptibility to β-lactams and cycloserine

Peptidoglycan precursor synthesis occurs in the cytoplasm and is inhibited at an early step by CYC. CYC is a dual inhibitor of alanine racemases (Alr and DadX), which synthesize D-Ala from L-Ala, and D-Ala-D-Ala ligase (Ddl), which synthesizes D-Ala-D-Ala (60). In *E. coli* and most Gram-negatives, peptidoglycan precursors contain a pentapeptide stem of L-Ala-D-Glu-*meso*-diaminopimelate-D-Ala-D-Ala. Glutamate racemase (MurI) is needed to convert L-Glu to D-Glu (61). Although CYC has poor inhibitory activity against K56-2 (MIC of 512 µg/mL), it is still useful as a chemical probe of biological function.

The targets of CYC are encoded by the *ddl*, *alr*, and *dadX* genes. While *ddl* is essential in K56-2 and not represented in our screen, we found that disruptions in *alr* and *dadX* caused susceptibility to CYC (Figure 5A). In *P. aeruginosa,* DadX is known to be more important for growth in nutrient-poor medium (62). As the BarSeq experiment was performed in rich LB medium, we observed that the fitness defect due to *dadX* disruption was less than that of *alr* and only the CRISPRi knockdown of *alr*, but not *dadX*, increased susceptibility to CYC (Figure 5B). Conversely, a disruption in *dadA*, encoding a catabolic D-amino acid dehydrogenase, resulted in moderately reduced susceptibility to CYC (Figure 5A). Knocking down *dadA* with CRISPRi increased the MIC of CYC by 2-fold (Figure 5B). Cells lacking DadA may have elevated pools of D-amino acids, including D-alanine, thus reducing dependence on the Alr and DadX alanine racemases.

**Figure 5.**
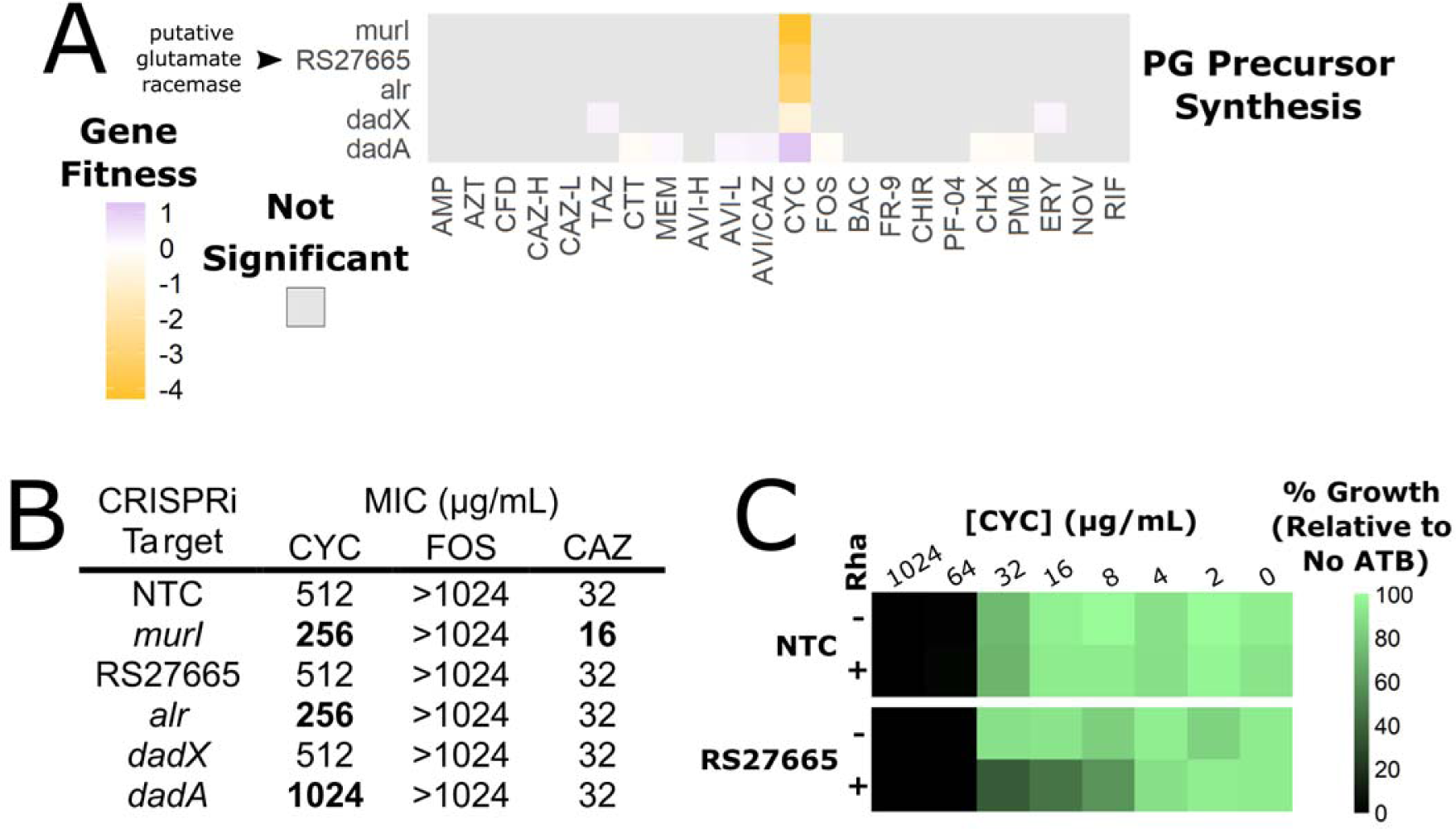
Disrupting cytoplasmic steps of peptidoglycan precursor synthesis increases cycloserine susceptibility. A) BarSeq gene fitness scores relative to the DMSO control of select genes associated with peptidoglycan precursor synthesis. Grey indicates the interactions were not significant (P > 0.05). B) MIC values of K56-2 CRISPRi mutants harbouring plasmids expressing a non-targeting sgRNA control (NTC) or a sgRNA targeting the indicated genes. Values shown are MIC values (medians of three biological replicates) in the presence of rhamnose induction of CRISPRi, with bold indicating at least a 2-fold change from the NTC. C) Cycloserine dose response of growth of a CRISPRi mutant in K562_RS27665 with or without 0.5% rhamnose. Values are averages of three biological replicates and normalized to the OD_600_ of growth without cycloserine.

Intriguingly, disruption of *murI* and K562_RS27665, encoding a putative fusion with GntR-type transcriptional regulator and pyridoxal phosphate (PLP)-dependent aminotransferase domains, also rendered the cells highly susceptible to CYC (Figure 5A). While CRISPRi silencing of only *murI* rendered a 2-fold reduction in the MIC of CYC (Figure 5B), silencing K562_RS27665 produced a subtle phenotype with reduced growth at subinhibitory CYC concentrations (Figure 5C). K562_RS27665 is homologous to the *E. coli* PatA putrescine aminotransferase and the uncharacterized *E. coli* proteins YdcR, YjiR, and YbdL. Interactions of CYC with D-glutamate biosynthesis have not been reported before. MurI is the only glutamate racemase in *E. coli* that synthesizes D-Glu for incorporation into peptidoglycan precursors; hence, *murI* is essential (63). Species of *Staphylococcus* and *Bacillus* encode glutamate racemase and a PLP-dependent D-amino acid transaminase capable of synthesizing D-glutamate using D-alanine and α-ketoglutarate as substrates (64, 65). With a score above 0.95, ProteInfer (66) confidently predicts that K562_RS27665 is associated with amino acid metabolic GO terms such as “L-aspartate: α-ketoglutarate aminotransferase activity” and “alanine catabolic process.” While the overall process of D-glutamate synthesis is essential, we suggest that because MurI and K562_RS27665 are nonessential in K56-2, they may be functionally redundant (67). Together, the chemical-genetic interactions with CYC suggest a delicate balance of D-amino acid metabolism required for peptidoglycan synthesis.

### Divisome-associated proteins are required for fitness when lateral and septal peptidoglycan synthesis is chemically inhibited

Peptidoglycan synthesis at the septum allows daughter-cell separation and is coordinated by the macromolecular complex termed the divisome (68). With the close association of divisome formation and peptidoglycan synthesis, we hypothesized that disruptions in cell division factors may increase susceptibility to cell envelope-targeting antibiotics. From the BarSeq experiment, we observed that disruption of *zapA, zapD,* and *ftsK* caused susceptibility to many β-lactams (Figure 6A). There was no significant effect of *zapE* disruption on fitness (Figure 6A); K56-2 lacks genes with strong similarity to *zapB* and *zapC*. In *E. coli*, these proteins all function at the divisome: FtsZ-ring associated proteins (Zaps) increase the cross-linking of FtsZ filaments, while FtsK aids divisome formation and chromosome segregation (68, 69). We found that CRISPRi knockdown of *zapA* increased susceptibility to AZT, CAZ, CFD, TAZ, and MEM (Figure 6B). In contrast, knockdown of *zapD* only increased susceptibility to AZT, CAZ, and MEM (Figure 6B). Neither mutant displayed a growth defect in the absence of antibiotics (Figure S3A). Depletion of ZapA and ZapD may increase susceptibility to β-lactams targeting septal PBPs (e.g. AZT) by reducing the efficiency of divisome formation, thus weakening the septal peptidoglycan matrix. In line with this, cell length significantly increased when *zapA* and *zapD* were silenced (Figure 6C and 6D). This effect was more pronounced in the presence of 0.25x MIC of AZT and MEM (Figure 6C and 6D). In the absence of rhamnose, there was no difference between the mutants and the non-targeting control mutant (Figure S5). Overall, these findings highlight the intrinsic interactions of cell division and peptidoglycan synthesis.

**Figure 6.**
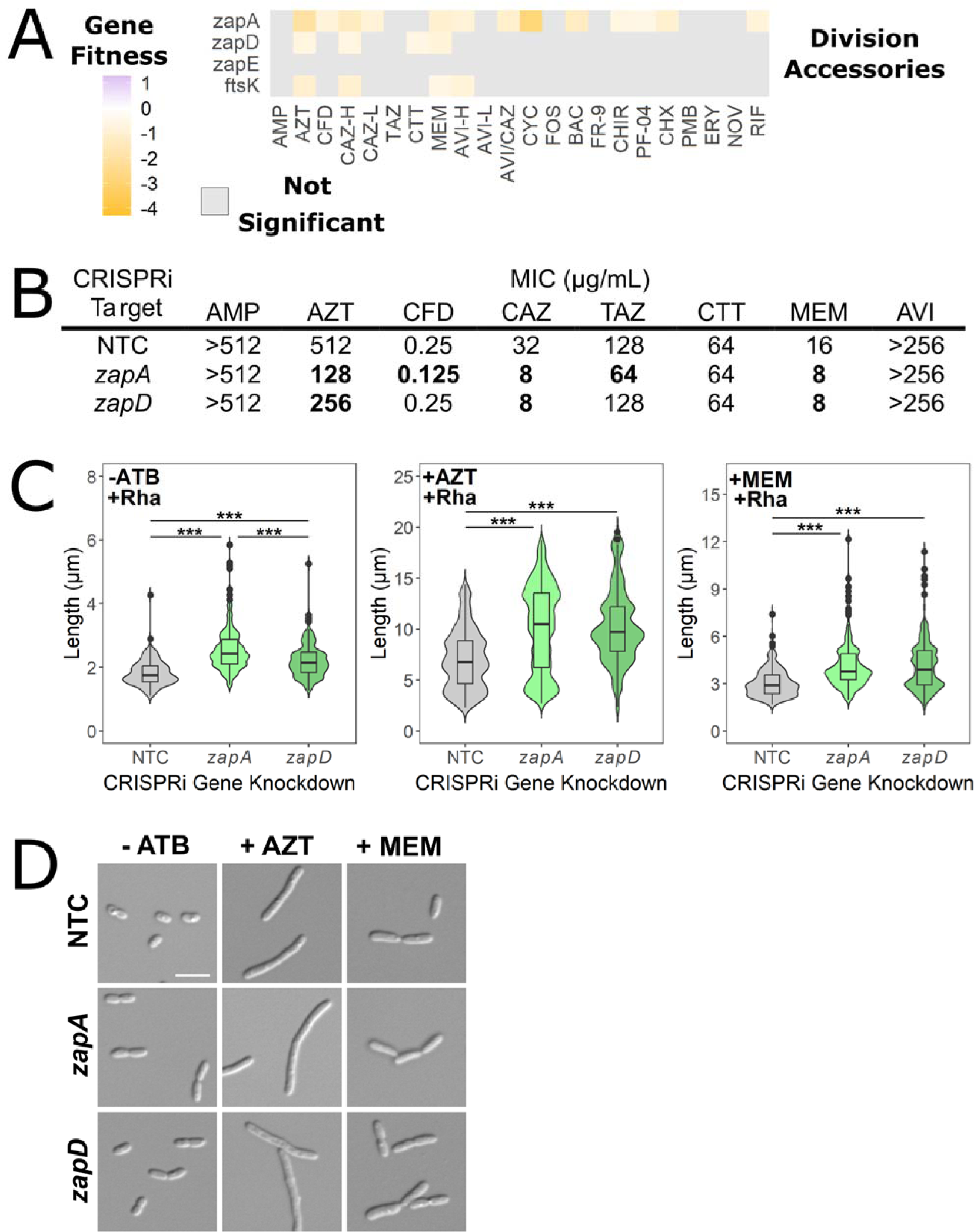
ZapA and ZapD are important for control of cell length and β-lactam susceptibility. A) BarSeq gene fitness scores relative to the DMSO control of cell division accessory genes. Grey indicates the interactions were not significant (P > 0.05). B) MIC values of K56-2 CRISPRi mutants silencing *zapA*, *zapD*, or a non-targeting control (NTC). Values shown are the medians of three biological replicates performed in the presence of rhamnose. C) Cell length distributions of CRISPRi mutants in *zapA* and *zapD* exposed to 0.25x the wild-type MIC of AZT (128 µg/mL) and MEM (4 µg/mL) for 90 minutes. In each condition, the length of 200 cells was recorded. Significance was determined by a Kruskal-Wall *H* test followed by Dunn’s *post hoc* test for pairwise differences. *** P < 0.0001. D) Representative micrographs of cells from Panel C. Scale bar = 5 µm.

## Discussion

The Gram-negative envelope is the first barrier to antibiotic activity and a substantial driver of antibiotic resistance. Here, we used previously reported data from a transposon screen in *B. cenocepacia* K56-2 exposed to an antibiotic panel, re-examining it in the context of cell envelope structural and functional components. Together with our recent report highlighting the roles of the Mla pathway, undecaprenyl phosphate recycling, β-lactamases, and TonB-dependent receptors (26), our present work furthers knowledge of how factors in the *Burkholderia* cell envelope contribute to envelope integrity and antibiotic resistance.

High levels of cationic antibiotic resistance in *B. cenocepacia* are suggested to stem from a “two-tier” model involving contributions from LPS and adaptive responses to cell envelope damage (31). We confirmed previous findings that disruptions in many genes along the O-antigen, LPS core, and lipid A modification pathways caused susceptibility to PMB (22, 32). We also observed that the same pathways are important for protection against CHX (Figure 1). Our work suggests that the O-antigen, LPS core, and Ara4N modification work together to hinder cationic antimicrobials from accessing their lipid membrane binding sites. The extent of resistance factors to cationic antibiotics, including the integration of the Ara4N modification into an essential network (67), suggests *Burkholderia* species were under strong and long-term environmental selection for antimicrobial peptide resistance.

Potentiators, also called adjuvants, are small molecules that synergize with antibiotics and generally lack growth-inhibitory activity on their own. Transposon mutagenesis experiments can identify genes that modulate antibiotic susceptibility *en masse*, some of which may be pursued as targets of potentiators. We recently used a transposon screen to link the potentiating activity of the β-lactamase inhibitor avibactam primarily to inhibition of the PenB carbapenemase in *B. cenocepacia* K56-2 (26). Here, we found that disruption of the DsbAB, but not the DsbCD, pathway substantially increased susceptibility to many antibiotics, including β-lactams and large-scaffold antibiotics. We linked susceptibility changes to reduced β-lactamase activity and increased outer membrane permeability (Figure 3). Notably, though, disruptions in DsbAB reduced susceptibility to cefiderocol, likely due to improper folding of the TonB-dependent receptors required to import this siderophore-cephalosporin conjugate (Table S3) (26). Another transposon screen by the Manoil group also found that DsbA and DsbB were important for resistance to meropenem in *A. baumannii* (70). In *E. coli*, the DsbAB pathway was recently linked to the stability of β-lactamases and mobile colistin resistance (MCR) enzymes (49). In *B. pseudomallei* and *B. cepacia*, the DsbAB pathway is important for antibiotic resistance and *in vivo* virulence (45, 47, 71). The pleiotropic effect of the DsbAB pathway makes it an attractive target for further investigation. Indeed, several inhibitors with *in vitro* and *in vivo* activity have been identified for DsbA (72, 73) and DsbB (74, 75), but none have yet been tested on *Burkholderia* species.

The peptidoglycan matrix is actively remodeled during cell growth and incorporates precursors from *de novo* synthesis and recycling. The balance of different precursors is tied into β-lactamase regulation; however, little is known about this process in *Burkholderia*. From the BarSeq screen in *B. cenocepacia* K56-2, we found that disruptions in each *dacB*, *slt*, *mltD*, *mltG*, and *ampG* increased β-lactam susceptibility. In contrast, disruption of *ampD* reduced β-lactam susceptibility (Figure 4). In *P. aeruginosa*, disruptions of *mltF*, *mltG*, *slt*, and *ampG* increase susceptibility to various β-lactams, often by reducing β-lactamase activity (76, 77). Additionally, disruption of *dacB* in *P. aeruginosa* greatly increased *ampC* expression and subsequently, β-lactam resistance (50). However, we found that disruption of *dacB* reduced susceptibility to MEM and AVI but not other β-lactams. Yet, it is unclear if these differences are due to changes in DacB function itself or how DacB is integrated into regulatory circuitry. Thus, we highlight areas requiring more work to elucidate mechanistic differences between other models and *B. cenocepacia* K56-2.

Steps in peptidoglycan recycling may be targets for potentiator development as their inactivation alters β-lactam susceptibility. The best-characterized inhibitor of this group is bulgecin A (78), a natural product inhibitor of transglycosylases Slt, MltD, and MltG in *P. aeruginosa* (79). In combination with β-lactams, bulgecin A rapidly induces midcell bulges and cell lysis (78). Potentiation of β-lactam activity has been observed in *E. coli, P. aeruginosa,* and *Neisseria* species (78–80). Other inhibitors active only *in vitro* include *N-*acetylglucosamine thiazoline (81) and thionine (82), which both target MltB (Slt35). Given the paucity of inhibitors, there is great potential in this area for the development of β-lactam potentiators.

The broad spectrum of bacterial efflux systems is an important consideration for antibiotic development and use. The RND family pumps in *B. cenocepacia* confer resistance to aminoglycosides, fluoroquinolones, trimethoprim, and chloramphenicol, among others (24, 40, 83). We found that NOV, AVI, and PF-04 are also substrates for the BpeAB-OprB efflux pump in *B. cenocepacia* K56-2 (Figure 2). BpeAB-OprB is a homologue of *P. aeruginosa* MexAB-OprM, known to efflux these compounds (84–86). Efflux of AVI is particularly concerning as this β-lactamase inhibitor is commonly used to restore β-lactam sensitivity to resistant Gram-negatives. Overexpression of efflux pumps is known to break the synergy of the AVI/CAZ combination in *P. aeruginosa* clinical isolates (86–88); however, there have been no reports of this effect in *Burkholderia* clinical isolates.

In summary, we report how chemical genetics can exploit antibiotics as specific probes to dissect contributions to cell envelope structure and function. The diverse panel of antibiotics we used revealed roles of the outer membrane, periplasm, and cytoplasm components to envelope integrity. Several of these functional annotations appear to be unique to *Burkholderia* (e.g. OpnA with NOV, LPS modifications with cationic peptides, DacB with β-lactams, and MurI/K562_RS27665 with CYC), highlighting important differences in *Burkholderia* physiology for further study.

## Methods and Materials

### Strains, medium preparation, and growth conditions

All strains used in this study (Table S5) were grown and maintained in LB-Lennox medium (Difco). All strains and mutants of *E. coli* and *B. cenocepacia* were grown at 37°C. *R. pickettii* K-288, *B. thailandensis* E264, *B. multivorans* ATCC 17616, *B. contaminans* FFH2055, *B. cepacia* ATCC 25416, *P. xenovorans* LB400, and *T. caryophylli* DSM 50341 were grown at 30°C.

All plasmids used in this study are listed in Table S6. Selective antibiotics were used at the following concentrations: trimethoprim (Sigma; 50 µg/mL for *E. coli* and 100 µg/mL for *Burkholderia* species), kanamycin (Fisher Scientific; 40 µg/mL for *E. coli*), gentamicin (Fisher Scientific; 50 µg/mL for *Burkholderia*), and tetracycline (Sigma; 20 µg/mL for *E. coli* and 100 µg/mL for *B. cenocepacia*).

### Molecular biology techniques

Previously reported methods were used for plasmid construction (overexpression and CRISPRi) and unmarked gene deletion in K56-2 (26, 41, 89). Briefly, primers with desired 5’ extensions (restriction sites or new sgRNA targeting regions) were ordered from IDT (Table S7). Q5 high-fidelity polymerase (NEB) with high-GC buffer was used to amplify DNA fragments for cloning. Fragments with restriction sites were digested with the appropriate restriction enzyme (NEB) and ligated with T4 ligase (NEB). *E. coli* DH5α was used to maintain plasmids. OneTaq polymerase with high-GC buffer (NEB) was used for colony PCR screening.

A previously designed Python script (90) was used to design sgRNAs next to the PAMs closest to the gene start. We used the same sgRNA stringency as reported before (26). Inverse PCR followed by blunt-end ligation was used to introduce new sgRNA targeting sites into pSCB2-sgRNAv2 as previously described (41, 91, 92).

Plasmids were typically introduced into Bcc species by triparental mating with the donor *E. coli* strain, the *E. coli* MM294/pRK2013 helper strain and the *Burkholderia* recipient (89). Exconjugants were selected with the appropriate antibiotics and 50 µg/mL gentamicin to select against *E. coli*. Successful plasmid mobilization or chromosomal changes were confirmed by colony PCR, as above.

The *wbiI* gene was deleted with double homologous recombination as per (93, 94) using a ∼900 bp upstream and downstream region fusion ordered as a single gBlock (IDT) (Table S8) with *Xma*I and *Xba*I restriction sites.

### Transposon mutagenesis, chemogenomics screening conditions, and data analysis

Construction of the barcoded transposon mutant library, exposure to the antibiotic panel and detection of individual mutant abundance were all performed in a previous publication (26). Briefly, a pool of ∼340,000 unique mutants was grown to exponential phase with 0.2% rhamnose (to allow outward gene expression and prevent transposon polar effects). In replicate 2mL culture at OD_600_ 0.15, the pools were exposed to individual antibiotics at concentrations equivalent to the IC_20_ – IC_30_ for 8 hours at 37°C, after which cells were harvested. A DMSO and Time 0 control were also prepared. Genomic DNA was extracted with the PureLink Genomic DNA Mini Kit (Invitrogen) and used as a template for barcode amplification in a 1-step PCR with Q5 polymerase with high-GC buffer (NEB). Indexed samples were sequenced on a NextSeq 550 in high-output mode (Donnelly Centre, Toronto, Canada) with reagent kit v2.5 and a 20% PhiX spike. The first 18 bases of sequencing were dark cycles to go past the zero-diversity flanking primer region. Barcode reads were linked to genomic insertion positions, and then mutant abundance and gene fitness in each condition were calculated using scripts from (95, 96).

For easier visualization, the gene fitness scores can be viewed in our interactive web application, which is available at https://cardonalab.shinyapps.io/bcc_interaction_viewer/.

### LPS analysis

The O-antigen and lipid A core were qualitatively analysed by silver staining as per (97), with the modifications reported in (26). Overnight cultures of the appropriate strains were diluted to OD_600_ 0.025 and grown to early exponential phase, then induced with 0.05% rhamnose for an additional 4 hours. Cells were harvested and lysed in a hot detergent buffer, followed by digestion with DNase I (Worthington Biochemical), then Proteinase K (ThermoFisher). Digested products were extracted in phenol (ThermoFisher) (supplemented with 0.1% β-mercaptoethanol and 0.2% 8-hydroxyquinoline) and washed with 10x volumes of ethyl ether. Gel loading dye was added to the aqueous phase in equal volume, run on a 14% Tricine SDS gel, and then stained with the Pierce Silver Stain Kit (ThermoFisher).

### Rhamnose dose-response curves

We determined how varying protein overexpression or CRISPRi knockdown affected growth after each mutant was created and before using the mutant in further physiological assays. A rhamnose gradient (starting at 1% for CRISPRi mutants and 0.2% for overexpression strains) was set up in 96-well format in LB medium. Each mutant was diluted to a final OD_600_ of 0.01 in 200 µL final volume in the gradient. Trimethoprim at 100 µg/mL was added to maintain selection. Growth with constant shaking at 37°C was monitored in a BioTek Synergy 2 plate reader.

### Antimicrobial susceptibility testing and interaction assays

Broth microdilution was used to assess susceptibility, and MIC values were interpreted per CLSI guidelines. Overnight cultures were used to prepare all inocula. Following 20 hours of static growth at either 30°C or 37°C, depending on the strain, growth was visually assessed.

Minor modifications were used when handling the CRISPRi mutants. Rhamnose at a final concentration of 0.5% was added to the wells to induce dCas9 expression. However, rhamnose was used at 0.05% for the mutant in *dsbB* as strong knockdown of this gene severely impacted growth (Figure S3A). Trimethoprim selection was not included to remove the possibility of antibiotic interactions.

Miniature checkerboard assays were prepared in CAMHB medium in 4×4 well format using 1/8 – 1/2 the MIC for each antimicrobial. Replicate checkerboard data was processed and interpreted with SynergyFinder2 (http://www.synergyfinder.org/#!/) (98, 99). Strong antagonistic interactions were noted when the Loewe Additivity and Bliss Independence scores were below - 10. On the other hand, when the scores were both above +10, the interactions were noted as strongly synergistic. Weak, synergistic or antagonistic interactions were noted when one score was above +10 or below −10 and the other was between −10 and +10. If both scores were between −10 and +10, no interaction was noted.

### NPN uptake assay

Outer membrane permeability was measured with an NPN uptake assay (33, 34), with modifications set out in (26). Briefly, overnight cultures of CRISPRi mutants were grown with rhamnose induction at 0.5%, diluted to OD_600_ 0.025, then grown for 8 hours with 0.5% rhamnose until OD_600_ ∼1.0. For the mutant in *dsbB*, 0.05% rhamnose was used to avoid a severe growth defect (Figure S3A). For gene deletion mutants, overnight cultures were diluted to OD_600_ 0.2 and grown until OD_600_ ∼1.0. Cells were then harvested and washed by centrifugation, then rested at room temperature in HEPES buffer with 10 mM NaN_3_ to depolarize for 30 minutes. NPN (Sigma) was then added to 40 µM final concentration, and the fluorescence was recorded in a BioTek Synergy 2 plate reader with filter sets: Excitation 360/40 nm and Emission 420/40 nm. Blank-corrected fluorescence values were normalized to the OD_600_ of each well.

### Nitrocefin hydrolysis assay of β-lactamase activity

This assay was performed as per (25, 100) with modifications reported in (26). Briefly, overnight cultures of CRISPRi mutants were grown with rhamnose induction at 0.5%, diluted to OD_600_ 0.1, then grown with 0.5% rhamnose until OD_600_ ∼2.0. For the mutant in *dsbB*, 0.05% rhamnose induction was used. Cells were harvested, resuspended in lysis buffer, freeze-thawed rapidly at −80°C, and then lysed by probe sonication. After centrifugation, the clarified supernatant was mixed with an equal volume of phosphate buffer with nitrocefin (final concentration of 100 µM nitrocefin (Sigma)). Sample absorbance over time was read at 485 nm on a BioTek Synergy 2 plate reader to calculate the rate of nitrocefin hydrolysis. This rate was normalized to the protein concentration of each sample, as determined by a BCA assay (Pierce), to calculate the specific activity.

### Bioinformatic screen for putative DSB system substrates

All protein-coding genes in *B. cenocepacia* K56-2 were analysed for predicted secretion signals by SignalP 5.0 with default settings (48). The number of cysteines was then counted in the predicted secreted proteins with the signal removed. Genes were annotated with eggnog-mapper v2 (101, 102). Functional enrichment in GO terms was calculated with GeneMerge 1.5 (103).

### Microscopy studies of zapA and zapD CRISPRi mutants

Overnight cultures of the mutants were grown with or without 0.5% rhamnose and 100 µg/mL trimethoprim. The mutants were then diluted to OD_600_ 0.025 and subcultured to exponential phase for 3 hours with or without 0.5% rhamnose and 100 µg/mL trimethoprim. The cultures were then exposed to 0.25x the wild-type K56-2 MIC of AZT (128 µg/mL) and MEM (4 µg/mL) for 90 minutes. An aliquot of cells was then washed by centrifugation and resuspended in Dulbecco’s PBS (Sigma) with 3.7% formaldehyde and 1% methanol. Cells were fixed at room temperature for 30 minutes. Unreacted formaldehyde was quenched by adding an equal volume of 0.5 M glycine. Cells were spotted on 1.5 % agarose pads and imaged by DIC microscopy with a Zeiss AxioImager. In each condition, the length of 200 cells was recorded. For differences between mutants, we performed a Kruskal-Wallis *H* test followed by Dunn’s *post hoc* test.

### RNA extraction and qRT-PCR analysis of gene expression

Assays were performed as previously reported (26, 41). Briefly, CRISPRi mutants were grown overnight with or without 0.5% rhamnose (0.05% for the *dsbB* mutant), then subcultured for 8 hr with the same amount of rhamnose at OD_600_ 0.01. Cells were harvested, and RNA was extracted with the PureLink RNA Mini kit (Invitrogen). RNA integrity was assessed by agarose gel electrophoresis. The ezDNase and SuperScript IV VILO RT (Invitrogen) were used for DNase treatment and cDNA synthesis, respectively. PowerTrack SYBR Green Mastermix (Invitrogen) was used for qRT-PCR with a StepOnePlus system (Applied Biosystems). Primer efficiency was assessed with cDNA serial dilutions (Table S9); only primers with 90-110% efficiency were used. Relative expression was quantified with the comparative C_T_ method (104) and normalized to the RNA polymerase sigma factor 70 housekeeping gene, *rpoD*.

### Data availability

The raw sequencing data generated in the previous report (26) was deposited in the NCBI Sequencing Read Archive (SRA) under BioProject ID PRJNA859150. The gene fitness scores from all antibiotic conditions are also available in the previous report (26).

## Acknowledgments

This work was financially supported by a Discovery Grant from the Natural Sciences and Engineering Research Council of Canada (NSERC), a Basic Research Grant from Cystic Fibrosis Canada, and a Research Project Grant from the Canadian Institutes of Health Research (CIHR) to STC; AMH was supported by a Vanier Canada Graduate Scholarship from CIHR.

## References

1. Zgurskaya HI, Lopez CA, Gnanakaran S. 2015. Permeability barrier of Gram-negative cell envelopes and approaches to bypass it. ACS InfectDis 1:512–522.

2. Murray CJ, Ikuta KS, Sharara F, Swetschinski L, Aguilar GR, Gray A, Han C, Bisignano C, Rao P, Wool E, Johnson SC, Browne AJ, Chipeta MG, Fell F, Hackett S, Haines-Woodhouse G, Hamadani BHK, Kumaran EAP, McManigal B, Agarwal R, Akech S, Albertson S, Amuasi J, Andrews J, Aravkin A, Ashley E, Bailey F, Baker S, Basnyat B, Bekker A, Bender R, Bethou A, Bielicki J, Boonkasidecha S, Bukosia J, Carvalheiro C, Castañeda-Orjuela C, Chansamouth V, Chaurasia S, Chiurchiù S, Chowdhury F, Cook AJ, Cooper B, Cressey TR, Criollo-Mora E, Cunningham M, Darboe S, Day NPJ, Luca MD, Dokova K, Dramowski A, Dunachie SJ, Eckmanns T, Eibach D, Emami A, Feasey N, Fisher-Pearson N, Forrest K, Garrett D, Gastmeier P, Giref AZ, Greer RC, Gupta V, Haller S, Haselbeck A, Hay SI, Holm M, Hopkins S, Iregbu KC, Jacobs J, Jarovsky D, Javanmardi F, Khorana M, Kissoon N, Kobeissi E, Kostyanev T, Krapp F, Krumkamp R, Kumar A, Kyu HH, Lim C, Limmathurotsakul D, Loftus MJ, Lunn M, Ma J, Mturi N, Munera-Huertas T, Musicha P, Mussi-Pinhata MM, Nakamura T, Nanavati R, Nangia S, Newton P, Ngoun C, Novotney A, Nwakanma D, Obiero CW, Olivas-Martinez A, Olliaro P, Ooko E, Ortiz-Brizuela E, Peleg AY, Perrone C, Plakkal N, Ponce-de-Leon A, Raad M, Ramdin T, Riddell A, Roberts T, Robotham JV, Roca A, Rudd KE, Russell N, Schnall J, Scott JAG, Shivamallappa M, Sifuentes-Osornio J, Steenkeste N, Stewardson AJ, Stoeva T, Tasak N, Thaiprakong A, Thwaites G, Turner C, Turner P, Doorn HR van, Velaphi S, Vongpradith A, Vu H, Walsh T, Waner S, Wangrangsimakul T, Wozniak T, Zheng P, Sartorius B, Lopez AD, Stergachis A, Moore C, Dolecek C, Naghavi M. 2022. Global burden of bacterial antimicrobial resistance in 2019: a systematic analysis. Lancet 399:629–655.

3. Masi M, Réfregiers M, Pos KM, Pagès J-M. 2017. Mechanisms of envelope permeability and antibiotic influx and efflux in Gram-negative bacteria. Nat Microbiol 2:17001.

4. Sun J, Rutherford ST, Silhavy TJ, Huang KC. 2022. Physical properties of the bacterial outer membrane. Nat Rev Microbiol 20:236–248.

5. Garde S, Chodisetti PK, Reddy M. 2021. Peptidoglycan: structure, synthesis, and regulation. EcoSal Plus 9.

6. Kamio Y, Nikaido H. 1976. The outer membrane of *Salmonella typhimurium*: accessibility of phospholipid head groups to phospholipase c and cyanogen bromide activated dextran in the external medium. Biochemistry 15:2561–2570.

7. Plésiat P, Nikaido H. 1992. Outer membranes of Gram-negative bacteria are permeable to steroid probes. Mol Microbiol 6:1323–1333.

8. Decad GM, Nikaido H. 1976. Outer membrane of Gram-negative bacteria. XII. Molecular-sieving function of cell wall. J Bacteriol 128:325–336.

9. Uehara T, Park JT. 2008. Peptidoglycan recycling. EcoSal Plus 3(1).

10. Blasco B, Pisabarro AG, de Pedro MA. 1988. Peptidoglycan biosynthesis in stationary-phase cells of *Escherichia coli*. J Bacteriol 170:5224–5228.

11. Payne DJ, Gwynn MN, Holmes DJ, Pompliano DL. 2007. Drugs for bad bugs: confronting the challenges of antibacterial discovery. Nat Rev Drug Discov 6:29–40.

12. Tommasi R, Brown DG, Walkup GK, Manchester JI, Miller AA. 2015. ESKAPEing the labyrinth of antibacterial discovery. Nat Rev Drug Discov 14:529–542.

13. Richter MF, Drown BS, Riley AP, Garcia A, Shirai T, Svec RL, Hergenrother PJ. 2017. Predictive compound accumulation rules yield a broad-spectrum antibiotic. Nature 545:299–304.

14. Leus IV, Weeks JW, Bonifay V, Shen Y, Yang L, Cooper CJ, Nath D, Duerfeldt AS, Smith JC, Parks JM, Rybenkov VV, Zgurskaya HI. 2022. Property space mapping of *Pseudomonas aeruginosa* permeability to small molecules. Sci Rep 12:8220.

15. Geddes EJ, Gugger MK, Garcia A, Chavez MG, Lee MR, Perlmutter SJ, Bieniossek C, Guasch L, Hergenrother PJ. 2023. Porin-independent accumulation in *Pseudomonas* enables antibiotic discovery. 7990. Nature 624:145–153.

16. Silver LL. 2016. A Gestalt approach to Gram-negative entry. Bioorg Med Chem 24:6379–6389.

17. Krishnamoorthy G, Leus IV, Weeks JW, Wolloscheck D, Rybenkov VV, Zgurskaya HI. 2017. Synergy between active efflux and outer membrane diffusion defines rules of antibiotic permeation into Gram-negative bacteria. mBio 8: e01172–17.

18. Leus IV, Adamiak J, Chandar B, Bonifay V, Zhao S, Walker SS, Squadroni B, Balibar CJ, Kinarivala N, Standke LC, Voss HU, Tan DS, Rybenkov VV, Zgurskaya HI. 2023. Functional diversity of Gram-negative permeability barriers reflected in antibacterial activities and intracellular accumulation of antibiotics. Antimicrob Agents Chemother 67:e01377–22.

19. Chmiel JF, Aksamit TR, Chotirmall SH, Dasenbrook EC, Elborn JS, LiPuma JJ, Ranganathan SC, Waters VJ, Ratjen FA. 2014. Antibiotic management of lung infections in cystic fibrosis. I. The microbiome, methicillin-resistant *Staphylococcus aureus*, Gram-negative bacteria, and multiple infections. Annals Am Thor Soc 11:1120–1129.

20. Isshiki Y, Kawahara K, Zähringer U. 1998. Isolation and characterisation of disodium (4-amino-4-deoxy-β-l-arabinopyranosyl)-(1-8)-(d-glycero-α-d-talo-oct-2-ulopyranosylonate)-(2-4)-(methyl 3-deoxy-d-manno-oct-2-ulopyranosid)onate from the lipopolysaccharide of *Burkholderia cepacia*. Carb Res 313:21–27.

21. Ortega XP, Cardona ST, Brown AR, Loutet SA, Flannagan RS, Campopiano DJ, Govan JR, Valvano MA. 2007. A putative gene cluster for aminoarabinose biosynthesis is essential for *Burkholderia cenocepacia* viability. J Bacteriol 189:3639–3644.

22. Hamad MA, Di Lorenzo F, Molinaro A, Valvano MA. 2012. Aminoarabinose is essential for lipopolysaccharide export and intrinsic antimicrobial peptide resistance in *Burkholderia cenocepacia*. Mol Microbiol 85:962–974.

23. Parr JT, Moore RA, Moore LV, Hancock RE. 1987. Role of porins in intrinsic antibiotic resistance of *Pseudomonas cepacia*. Antimicrob Agents Chemother 31:121–123.

24. Scoffone VC, Trespidi G, Barbieri G, Irudal S, Perrin E, Buroni S. 2021. Role of RND efflux pumps in drug resistance of cystic fibrosis pathogens. Antibiotics 10:863.

25. Zeiser ET, Becka SA, Wilson BM, Barnes MD, LiPuma JJ, Papp-Wallace KM. 2019. “Switching partners”: piperacillin-avibactam is a highly potent combination against multidrug-resistant *Burkholderia cepacia* complex and *Burkholderia gladioli* cystic fibrosis isolates. J Clin Microbiol 57:e00181–19.

26. Hogan AM, Rahman ASMZ, Motnenko A, Natarajan A, Maydaniuk DT, León B, Batun Z, Palacios A, Bosch A, Cardona ST. 2023. Profiling cell envelope-antibiotic interactions reveals vulnerabilities to β-lactams in a multidrug-resistant bacterium. Nat Commun 14:4815.

27. Darling P, Chan M, Cox AD, Sokol PA. 1998. Siderophore production by cystic fibrosis isolates of *Burkholderia cepacia*. Infect Immun 66:874–877.

28. Sabnis A, Hagart KL, Klöckner A, Becce M, Evans LE, Furniss RCD, Mavridou DA, Murphy R, Stevens MM, Davies JC, Larrouy-Maumus GJ, Clarke TB, Edwards AM. 2021. Colistin kills bacteria by targeting lipopolysaccharide in the cytoplasmic membrane. eLife 10:e65836.

29. Barrett-Bee K, Newboult L, Edwards S. 1994. The membrane destabilising action of the antibacterial agent chlorhexidine. FEMS Microbiol Lett 119:249–253.

30. Simpson BW, Trent MS. 2019. Pushing the envelope: LPS modifications and their consequences. Nat Rev Microbiol 17:403–416.

31. Loutet SA, Mussen LE, Flannagan RS, Valvano MA. 2011. A two-tier model of polymyxin B resistance in *Burkholderia cenocepacia*. Environ Microbiol Rep 3:278–285.

32. Ortega X, Silipo A, Saldías MS, Bates CC, Molinaro A, Valvano MA. 2009. Biosynthesis and structure of the *Burkholderia cenocepacia* K56-2 lipopolysaccharide core oligosaccharide: truncation of the core oligosaccharide leads to increased binding and sensitivity to polymyxin B. J Biol Chem 284:21738–21751.

33. Loh B, Grant C, Hancock RE. 1984. Use of the fluorescent probe 1-N-phenylnaphthylamine to study the interactions of aminoglycoside antibiotics with the outer membrane of *Pseudomonas aeruginosa*. Antimicrob Agents Chemother 26:546–551.

34. Malott RJ, Steen-Kinnaird BR, Lee TD, Speert DP. 2012. Identification of hopanoid biosynthesis genes involved in polymyxin resistance in *Burkholderia multivorans*. Antimicrob Agents Chemother 56:464–471.

35. Gunn JS, Lim KB, Krueger J, Kim K, Guo L, Hackett M, Miller SI. 1998. PmrA–PmrB-regulated genes necessary for 4-aminoarabinose lipid A modification and polymyxin resistance. Mol Microbiol 27:1171–1182.

36. Trent MS, Ribeiro AA, Doerrler WT, Lin S, Cotter RJ, Raetz CR. 2001. Accumulation of a polyisoprene-linked amino sugar in polymyxin-resistant *Salmonella typhimurium* and *Escherichia coli*: Structural characterization and transfer to lipid A in the periplasm. J Biol Chem 276:43132–43144.

37. Panta PR, Kumar S, Stafford CF, Billiot CE, Douglass MV, Herrera CM, Trent MS, Doerrler WT. 2019. A DedA family membrane protein is required for *Burkholderia thailandensis* colistin resistance. Front Microbiol 10.

38. Sit B, Srisuknimit V, Bueno E, Zingl FG, Hullahalli K, Cava F, Waldor MK. 2022. Undecaprenyl phosphate translocases confer conditional microbial fitness. Nature 613:721.

39. Roney IJ, Rudner DZ. 2023. Two broadly conserved families of polyprenyl-phosphate transporters. 7945. Nature 613:729–734.

40. Bazzini S, Udine C, Sass A, Pasca MR, Longo F, Emiliani G, Fondi M, Perrin E, Decorosi F, Viti C, Giovannetti L, Leoni L, Fani R, Riccardi G, Mahenthiralingam E, Buroni S. 2011. Deciphering the role of RND efflux transporters in *Burkholderia cenocepacia*. PLoS ONE 6:e18902.

41. Hogan AM, Rahman ASMZ, Lightly TJ, Cardona ST. 2019. A broad-host-range CRISPRi toolkit for silencing gene expression in *Burkholderia*. ACS Synth Biol 8:2372–2384.

42. Choi U, Lee C-R. 2019. Distinct roles of outer membrane porins in antibiotic resistance and membrane integrity in *Escherichia coli*. Front Microbiol 10.

43. Landeta C, Boyd D, Beckwith J. 2018. Disulfide bond formation in prokaryotes. Nat Microbiol 3:270–280.

44. Abe M, Nakazawa T. 1996. The *dsbB* gene product is required for protease production by *Burkholderia cepacia*. Infect Immun 64:4378–4380.

45. Hayashi S, Abe M, Kimoto M, Furukawa S, Nakazawa T. 2000. The DsbA-DsbB disulfide bond formation system of *Burkholderia cepacia* is involved in the production of protease and alkaline phosphatase, motility, metal resistance, and multi-drug resistance. Microbiol Immunol 44:41–50.

46. Dutton RJ, Boyd D, Berkmen M, Beckwith J. 2008. Bacterial species exhibit diversity in their mechanisms and capacity for protein disulfide bond formation. PNAS 105:11933–11938.

47. Vezina B, Petit GA, Martin JL, Halili MA. 2020. Prediction of Burkholderia pseudomallei DsbA substrates identifies potential virulence factors and vaccine targets. PLoS One 15:e0241306.

48. Almagro Armenteros JJ, Tsirigos KD, Sønderby CK, Petersen TN, Winther O, Brunak S, von Heijne G, Nielsen H. 2019. SignalP 5.0 improves signal peptide predictions using deep neural networks. Nat Biotechnol 37:420–423.

49. Furniss RCD, Kaderabkova N, Barker D, Bernal P, Maslova E, Antwi AA, McNeil HE, Pugh HL, Dortet L, Blair JM, Larrouy-Maumus G, McCarthy RR, Gonzalez D, Mavridou DA. 2022. Breaking antimicrobial resistance by disrupting extracytoplasmic protein folding. eLife 11:e57974.

50. Moya B, Dötsch A, Juan C, Blázquez J, Zamorano L, Haussler S, Oliver A. 2009. β-lactam resistance response triggered by inactivation of a nonessential penicillin-binding protein. PLoS Path 5:e1000353.

51. Liu C, Li C, Chen Y, Hao H, Liang J, Duan R, Guo Z, Zhang J, Zhao Z, Jing H, Wang X, Shao S. 2017. Role of low-molecular-mass penicillin-binding proteins, NagZ and AmpR in AmpC β-lactamase regulation of *Yersinia enterocolitica*. Front Cell Infect Microbiol 7.

52. Tayler AE, Ayala JA, Niumsup P, Westphal K, Baker JA, Zhang L, Walsh TR, Wiedemann B, Bennett PM, Avison MB. 2010. Induction of beta-lactamase production in *Aeromonas hydrophila* is responsive to beta-lactam-mediated changes in peptidoglycan composition. Microbiology (Reading) 156:2327–2335.

53. Dik DA, Fisher JF, Mobashery S. 2018. Cell-wall recycling of the Gram-negative bacteria and the nexus to antibiotic resistance. Chem Rev 118:5952–5984.

54. Weaver AI, Alvarez L, Rosch KM, Ahmed A, Wang GS, van Nieuwenhze MS, Cava F, Dörr T. 2022. Lytic transglycosylases mitigate periplasmic crowding by degrading soluble cell wall turnover products. eLife 11:e73178.

55. Jacobs C, Huang LJ, Bartowsky E, Normark S, Park JT. 1994. Bacterial cell wall recycling provides cytosolic muropeptides as effectors for beta-lactamase induction. EMBO J 13:4684–4694.

56. Jacobs C, Joris B, Jamin M, Klarsov K, van Beeumen J, Mengin-Lecreulx D, van Heijenoort J, Park JT, Normark S, Frère J-M. 1995. AmpD, essential for both β-lactamase regulation and cell wall recycling, is a novel cytosolic N-acetylmuramyl-L-alanine amidase. Mol Microbiol 15:553–559.

57. Bartowsky E, Normark S. 1991. Purification and mutant analysis of *Citrobacter freundii* AmpR, the regulator for chromosomal AmpC beta-lactamase. Mol Microbiol 5:1715–1725.

58. Balcewich MD, Reeve TM, Orlikow EA, Donald LJ, Vocadlo DJ, Mark BL. 2010. Crystal structure of the AmpR effector binding domain provides insight into the molecular regulation of inducible AmpC beta-lactamase. J Mol Biol 400:998–1010.

59. Hwang J, Kim HS. 2015. Cell wall recycling-linked coregulation of AmpC and PenB β-lactamases through *ampD* mutations in *Burkholderia cenocepacia*. Antimicrob Agents Chemother 59:7602–7610.

60. Walsh CT. 1989. Enzymes in the D-alanine branch of bacterial cell wall peptidoglycan assembly. J Biol Chem 264:2393–2396.

61. Barreteau H, Kovač A, Boniface A, Sova M, Gobec S, Blanot D. 2008. Cytoplasmic steps of peptidoglycan biosynthesis. FEMS Microbiol Rev 32:168–207.

62. He W, Li C, Lu C-D. 2011. Regulation and characterization of the *dadRAX* locus for d-amino acid catabolism in *Pseudomonas aeruginosa* PAO1. J Bacteriol 193:2107–2115.

63. Doublet P, van Heijenoort J, Bohin JP, Mengin-Lecreulx D. 1993. The *murI* gene of *Escherichia coli* is an essential gene that encodes a glutamate racemase activity. J Bacteriol 175:2970–2979.

64. Pucci MJ, Thanassi JA, Ho HT, Falk PJ, Dougherty TJ. 1995. *Staphylococcus haemolyticus* contains two D-glutamic acid biosynthetic activities, a glutamate racemase and a D-amino acid transaminase. J Bacteriol 177:336–342.

65. Fotheringham IG, Bledig SA, Taylor PP. 1998. Characterization of the genes encoding d-amino acid transaminase and glutamate racemase, two d-glutamate biosynthetic enzymes of *Bacillus sphaericus* ATCC 10208. J Bacteriol 180:4319–4323.

66. Sanderson T, Bileschi ML, Belanger D, Colwell LJ. 2023. ProteInfer: deep networks for protein functional inference. eLife 12:e80942.

67. Gislason AS, Turner K, Domaratzki M, Cardona ST. 2017. Comparative analysis of the *Burkholderia cenocepacia* K56-2 essential genome reveals cell envelope functions that are uniquely required for survival in species of the genus *Burkholderia*. Microb Genom 3:e000140.

68. Du S, Lutkenhaus J. 2019. At the heart of bacterial cytokinesis: the Z ring. Trends Microbiol 27:781–791.

69. Aussel L, Barre F-X, Aroyo M, Stasiak A, Stasiak AZ, Sherratt D. 2002. FtsK is a DNA motor protein that activates chromosome dimer resolution by switching the catalytic state of the XerC and XerD recombinases. Cell 108:195–205.

70. Bailey J, Gallagher L, Barker WT, Hubble VB, Gasper J, Melander C, Manoil C. 2022. Genetic dissection of antibiotic adjuvant activity. mBio e0308421.

71. McMahon RM, Ireland PM, Sarovich DS, Petit G, Jenkins CH, Sarkar-Tyson M, Currie BJ, Martin JL. 2018. Virulence of the melioidosis pathogen *Burkholderia pseudomallei* requires the oxidoreductase membrane protein DsbB. Infect Immun 86:e00938–17.

72. Duprez W, Premkumar L, Halili MA, Lindahl F, Reid RC, Fairlie DP, Martin JL. 2015. Peptide Inhibitors of the *Escherichia coli* DsbA oxidative machinery essential for bacterial virulence. J Med Chem 58:577–587.

73. Adams LA, Sharma P, Mohanty B, Ilyichova OV, Mulcair MD, Williams ML, Gleeson EC, Totsika M, Doak BC, Caria S, Rimmer K, Horne J, Shouldice SR, Vazirani M, Headey SJ, Plumb BR, Martin JL, Heras B, Simpson JS, Scanlon MJ. 2015. Application of fragment-based screening to the design of inhibitors of *Escherichia coli* DsbA. Angew Chem Int Ed Engl 54:2179–2184.

74. Landeta C, Blazyk JL, Hatahet F, Meehan BM, Eser M, Myrick A, Bronstain L, Minami S, Arnold H, Ke N, Rubin EJ, Furie BC, Furie B, Beckwith J, Dutton R, Boyd D. 2015. Compounds targeting disulfide bond forming enzyme DsbB of Gram-negative bacteria. Nat Chem Biol 11:292–298.

75. Landeta C, McPartland L, Tran NQ, Meehan BM, Zhang Y, Tanweer Z, Wakabayashi S, Rock J, Kim T, Balasubramanian D, Audette R, Toosky M, Pinkham J, Rubin EJ, Lory S, Pier G, Boyd D, Beckwith J. 2019. Inhibition of *Pseudomonas aeruginosa* and *Mycobacterium tuberculosis* disulfide bond forming enzymes. Mol Microbiol 111:918–937.

76. Cavallari JF, Lamers RP, Scheurwater EM, Matos AL, Burrows LL. 2013. Changes to its peptidoglycan-remodeling enzyme repertoire modulate β-lactam resistance in *Pseudomonas aeruginosa*. Antimicrob Agents Chemother 57:3078–3084.

77. Sonnabend MS, Klein K, Beier S, Angelov A, Kluj R, Mayer C, Groß C, Hofmeister K, Beuttner A, Willmann M, Peter S, Oberhettinger P, Schmidt A, Autenrieth IB, Schütz M, Bohn E. 2020. Identification of drug resistance determinants in a clinical isolate of *Pseudomonas aeruginosa* by high-density transposon mutagenesis. Antimicrob Agents Chemother 64:e01771–19.

78. Imada A, Kintaka K, Nakao M, Shinagawa S. 1982. Bulgecin, a bacterial metabolite which in concert with beta-lactam antibiotics causes bulge formation. J Antibiot (Tokyo) 35:1400–1403.

79. Dik DA, Madukoma CS, Tomoshige S, Kim C, Lastochkin E, Boggess WC, Fisher JF, Shrout JD, Mobashery S. 2019. Slt, MltD, and MltG of *Pseudomonas aeruginosa* as targets of bulgecin A in potentiation of β-lactam antibiotics. ACS Chem Biol 14:296–303.

80. Williams AH, Wheeler R, Thiriau C, Haouz A, Taha M-K, Boneca IG. 2017. Bulgecin A: the key to a broad-spectrum inhibitor that targets lytic transglycosylases. Antibiotics 6:8.

81. Reid CW, Blackburn NT, Legaree BA, Auzanneau F-I, Clarke AJ. 2004. Inhibition of membrane-bound lytic transglycosylase B by NAG-thiazoline. FEBS Lett 574:73–79.

82. Mezoughi AB, Costanzo CM, Parker GM, Behiry EM, Scott A, Wood AC, Adams SE, Sessions RB, Loveridge EJ. 2021. The lysozyme inhibitor thionine acetate is also an inhibitor of the soluble lytic transglycosylase Slt35 from *Escherichia coli*. Molecules 26:4189.

83. Buroni S, Pasca MR, Flannagan RS, Bazzini S, Milano A, Bertani I, Venturi V, Valvano MA, Riccardi G. 2009. Assessment of three resistance-nodulation-cell division drug efflux transporters of *Burkholderia cenocepacia* in intrinsic antibiotic resistance. BMC Microbiol 9:200.

84. Srikumar R, Kon T, Gotoh N, Poole K. 1998. Expression of *Pseudomonas aeruginosa* multidrug efflux pumps MexA-MexB-OprM and MexC-MexD-OprJ in a multidrug-sensitive *Escherichia coli* strain. Antimicrob Agents Chemother 42:65–71.

85. Caughlan RE, Jones AK, Delucia AM, Woods AL, Xie L, Ma B, Barnes SW, Walker JR, Sprague ER, Yang X, Dean CR. 2012. Mechanisms decreasing in vitro susceptibility to the LpxC inhibitor CHIR-090 in the Gram-negative pathogen *Pseudomonas aeruginosa*. Antimicrob Agents Chemother 56:17–27.

86. Torrens G, Cabot G, Ocampo-Sosa AA, Conejo MC, Zamorano L, Navarro F, Pascual Á, Martínez-Martínez L, Oliver A. 2016. Activity of ceftazidime-avibactam against clinical and isogenic laboratory *Pseudomonas aeruginosa* isolates expressing combinations of most relevant β-lactam resistance mechanisms. Antimicrob Agents Chemother 60:6407–6410.

87. Chalhoub H, Sáenz Y, Nichols WW, Tulkens PM, Van Bambeke F. 2018. Loss of activity of ceftazidime-avibactam due to MexAB-OprM efflux and overproduction of AmpC cephalosporinase in *Pseudomonas aeruginosa* isolated from patients suffering from cystic fibrosis. Int J Antimicrob Agents 52:697–701.

88. Castanheira M, Doyle TB, Smith CJ, Mendes RE, Sader HS. 2019. Combination of MexAB-OprM overexpression and mutations in efflux regulators, PBPs and chaperone proteins is responsible for ceftazidime/avibactam resistance in *Pseudomonas aeruginosa* clinical isolates from US hospitals. J Antimicrob Chemother 74:2588–2595.

89. Hogan AM, Scoffone VC, Makarov V, Gislason AS, Tesfu H, Stietz MS, Brassinga AKC, Domaratzki M, Li X, Azzalin A, Biggiogera M, Riabova O, Monakhova N, Chiarelli LR, Riccardi G, Buroni S, Cardona ST. 2018. Competitive fitness of essential gene knockdowns reveals a broad-spectrum antibacterial inhibitor of the cell division protein FtsZ. Antimicrob Agents Chemother 62:e01231–18.

90. van Gestel J, Hawkins JS, Todor H, Gross CA. 2021. Computational pipeline for designing guide RNAs for mismatch-CRISPRi. STAR Protocols 2:100521.

91. Larson MH, Gilbert LA, Wang X, Lim WA, Weissman JS, Qi LS. 2013. CRISPR interference (CRISPRi) for sequence-specific control of gene expression. Nat Protoc 8:2180–2196.

92. Qi LS, Larson MH, Gilbert LA, Doudna JA, Weissman JS, Arkin AP, Lim WA. 2013. Repurposing CRISPR as an RNA-guided platform for sequence-specific control of gene expression. Cell 152:1173–1183.

93. Flannagan RS, Linn T, Valvano MA. 2008. A system for the construction of targeted unmarked gene deletions in the genus *Burkholderia*. Environ Microbiol 10:1652–1660.

94. Hamad MA, Skeldon AM, Valvano MA. 2010. Construction of aminoglycoside-sensitive *Burkholderia cenocepacia* strains for use in studies of intracellular bacteria with the gentamicin protection assay. Appl Environ Microbiol 76:3170–3176.

95. Wetmore KM, Price MN, Waters RJ, Lamson JS, He J, Hoover CA, Blow MJ, Bristow J, Butland G, Arkin AP, Deutschbauer A. 2015. Rapid quantification of mutant fitness in diverse bacteria by sequencing randomly barcoded transposons. mBio 6:e00306–00315.

96. Morin M, Pierce EC, Dutton RJ. 2018. Changes in the genetic requirements for microbial interactions with increasing community complexity. eLife 7:e37072.

97. Marolda CL, Lahiry P, Vinés E, Saldías S, Valvano MA. 2006. Micromethods for the characterization of lipid A-core and O-antigen lipopolysaccharide. Methods Mol Biol 347:237–252.

98. Ianevski A, Giri AK, Aittokallio T. 2020. SynergyFinder 2.0: visual analytics of multi-drug combination synergies. Nucleic Acids Res 48:W488–W493.

99. Zheng S, Wang W, Aldahdooh J, Malyutina A, Shadbahr T, Tanoli Z, Pessia A, Tang J. 2022. SynergyFinder Plus: Toward better interpretation and annotation of drug combination screening datasets. Genom Proteom Bioinform S1672-0229(22)00008–0.

100. Alvarez-Ortega C, Wiegand I, Olivares J, Hancock REW, Martínez JL. 2010. Genetic determinants involved in the susceptibility of *Pseudomonas aeruginosa* to β-lactam antibiotics. Antimicrob Agents Chemother 54:4159–4167.

101. Huerta-Cepas J, Szklarczyk D, Heller D, Hernández-Plaza A, Forslund SK, Cook H, Mende DR, Letunic I, Rattei T, Jensen LJ, von Mering C, Bork P. 2019. eggNOG 5.0: a hierarchical, functionally and phylogenetically annotated orthology resource based on 5090 organisms and 2502 viruses. Nucleic Acids Res 47:D309–D314.

102. Cantalapiedra CP, Hernández-Plaza A, Letunic I, Bork P, Huerta-Cepas J. 2021. eggNOG-mapper v2: Functional annotation, orthology assignments, and domain prediction at the metagenomic scale. Mol Biol Evol 38: 5825–5829.

103. Castillo-Davis CI, Hartl DL. 2003. GeneMerge--post-genomic analysis, data mining, and hypothesis testing. Bioinformatics 19:891–892.

104. Schmittgen TD, Livak KJ. 2008. Analyzing real-time PCR data by the comparative C(T) method. Nat Protoc 3:1101–1108.

